# Fundamental contribution and host range determination of ANP32 protein family in influenza A virus polymerase activity

**DOI:** 10.1101/529412

**Authors:** Haili Zhang, Zhenyu Zhang, Yujie Wang, Meiyue Wang, Xuefeng Wang, Xiang Zhang, Shuang Ji, Cheng Du, Hualan Chen, Xiaojun Wang

## Abstract

The polymerase of the influenza virus is part of the key machinery necessary for viral replication. However, the avian influenza virus polymerase is restricted in mammalian cells. The cellular protein ANP32A has been recently found to interact with viral polymerase, and to both influence polymerase activity and interspecies restriction. Here we report that either ANP32A or ANP32B is indispensable for influenza A virus RNA replication. The contribution of ANP32B is equal to that of ANP32A, and together they play a fundamental role in the activity of mammalian influenza A virus polymerase, while neither human ANP32A nor ANP32B support the activity of avian viral polymerase. Interestingly, we found that avian ANP32B was naturally inactive, leaving ANP32A alone to support viral replication. Two amino acid mutations at sites 129-130 in chicken ANP32B lead to the loss of support of viral replication and weak interaction with the viral polymerase complex, and these amino acids are also crucial in the maintenance of viral polymerase activity in other ANP32 proteins. Our findings strongly support ANP32A&B as key factors for both virus replication and adaption.

**IMPORTANCE:** The key host factors involved in the influenza A viral the polymerase activity and RNA replication remain largely unknown. Here we provide evidence that ANP32A and ANP32B from different species are powerful factors in the maintenance of viral polymerase activity. Human ANP32A and ANP32B contribute equally to support human influenza virus RNA replication. However, unlike avian ANP32A, the avian ANP32B is evolutionarily non-functional in supporting viral replication because of a 129-130 site mutation. The 129-130 site plays an important role in ANP32A/B and viral polymerase interaction, therefore determine viral replication, suggesting a novel interface as a potential target for the development of anti-influenza strategies.

## INTRODUCTION

Virus transmission from its natural host species to a different species reflects mechanisms of molecular restriction, evolution, and adaptation. Birds harbor most of the influenza A viruses, which are typically replication-restricted in human hosts because of receptor incompatibility and limited viral polymerase activity in human cells. Although many host factors have been reported to be involved in viral replication (1-8), the key mechanisms that determine viral polymerase activity and host range are poorly understood.

Influenza A viral ribonucleoprotein (vRNP), the viral minimum replicon, comprises the viral genome, the heterotrimeric RNA polymerase PB1, PB2, PA, and the nucleoprotein (NP), and carries out viral RNA transcription and replication in infected cells. Avian influenza A polymerases have very limited activity in mammalian cells, indicating an unknown host-specific restriction mechanism that directly affects viral RNA replication. The adaptation of avian viruses to mammals, such as occurred with the H5N1 and H7N9 avian viruses, occurs along with substitutions on the viral polymerase, mainly on the PB2 subunit (E627K and other signatures), which enhance viral replication (9-22). Many host factors have been reported to interact with the vRNP complex to help viral replication (7, 23-33). Of these proteins, chicken acidic nuclear phosphoprotein 32 family member A (chANP32A) has been reported to specifically promote avian influenza replication due to the 33 amino acids insert (8). This 33 amino acid insert includes a hydrophobic SUMO interaction motif, which connects to host SUMOylation to partially contribute to the promotion of avian viral polymerase activity (34). Furthermore, ANP32A from birds has splicing variants without SUMO interaction motif also support viral replication and polymerase adaption (35). Interestingly, a previous study showed that ANP32A&B (acidic nuclear phosphoprotein 32 family, members A&B) have been found to regulate viral RNA synthesis (7). These studies suggest that ANP32A plays an important role in viral replication.

The acidic (leucine-rich) nuclear phosphoprotein 32kDa (ANP32) family comprises several members, including ANP32A, ANP32B, and ANP32E, which have various functions in the regulation of gene expression, intracellular transportation, and cell death (36). Although ANP32A is considered to be an important co-factor of the influenza virus polymerase and to influence the viral host range, the roles of different ANP32 members in viral replication, and the extent to which the proteins are involved in the activity of the polymerase remain unclear. In this study, by using CRISPR/Cas9 knockout screening, we found that ANP32A and ANP32B play fundamental roles in the facilitation of influenza A viral RNA synthesis, and that both ANP32A and ANP32B contribute equally. The mammalian ANP32 proteins give no or only limited support to the avian virus polymerase. Human ANP32A&B do not support the replication of the avian influenza virus, but this restriction can be overcome by E627K substitution in the viral PB2 protein. Furthermore, we found that chicken ANP32B has no effect on polymerase activity, which can be ascribed to mutations in two key amino acid residues of ANP32A&B that determine their activity in supporting viral RNA replication. Together, these data reveal fundamental roles for ANP32A and ANP32B in supporting influenza virus A polymerase activity as well as a site key for their function, and show that both ANP32A and B determine viral polymerase adaptation and host range.

## RESULTS

### HuANP32A&B Are Critical Host Factors that Determine Viral Polymerase Activity and Virus Replication

Previous studies have reported that several host factors, including BUB3, CLTC, CYC1, NIBP, ZC3H15, C14orf173, CTNNB1, ANP32A, ANP32B, SUPT5H, HTATSF1, and DDX17, interact with influenza viral polymerase and that some of these factors have an effect on viral polymerase activity (4, 23, 37). All these observations were based on the gene transient knockdown technique, meaning that the results may vary because of the different knockdown efficiency of target genes. Using a CRISPR/Cas9 system, however, allows rapid knockout of certain genes and accurate evaluation of target proteins. To identify the critical roles of the above-mentioned host factors in influenza viral replication, we used a CRISPR/Cas9 system to establish a series of knockout 293T cell lines, and a model viral-like luciferase RNA was expressed together with viral polymerase PB1, PB2, PA, and NP, to determine the polymerase activity in these cell lines. Firstly, we tested the polymerase activity of 2009 pandemic H1N1 virus A/Sichuan/01/2009(H1N1_SC09_) (38) on these knockout 293T cells, and found that individual knockout of NIBP, ZC3H15, or DDX17 results in a 2-4 fold decrease in viral polymerase activity, in which is consistent with previous results (23, 37). However, in none of the other single gene knockout cells was viral polymerase activity blocked (Fig. 1A). We did not observe any reduction of viral polymerase activity in ANP32A or ANP32B knockout cells, which was a surprising result as ANP32A and ANP32B were reported to be important host factors supporting viral polymerase activity (7, 8, 34, 35). Since ANP32A and ANP32B have high similarity in both structure and known functions, we predicted that in single knockout cell lines of ANP32A or ANP32B, the presence of either one of these two proteins could support viral replication in the absence of the other. We then developed an ANP32A and ANP32B double knockout cell line to confirm this hypothesis. In human ANP32A (huANP32A) knockout 293T cells (AKO cells), and human ANP32B (huANP32B) knockout 293T cells (BKO cells), the viral polymerases have similar activities to wild-type 293T cells, but when both ANP32A and ANP32B were knocked out (DKO) (Fig. S1 in the supplemental material), the polymerase activity was abolished (about 10000-fold reduction) (Fig 1B), although the polymerase was expressed equally in the WT, AKO, BKO, and DKO cells (Fig. 1B). We then tested the polymerase activities of 2013 China H7N9 human isolate A/Anhui/01/2013 (H7N9_AH13_) (39), and H1N1 virus A/WSN/1933 (WSN) on AKO, BKO and DKO cell lines, and found that the viral polymerase complex lost all activity in the DKO cells, but not in the AKO or BKO cells (Fig. 1C and 1D). We further confirmed the effect of huANP32A&B on viral infectivity by infecting 293T cells and DKO cells with WSN virus. In DKO cells, but not AKO or BKO cells, the infectivity of WSN decreased about 100000-fold (Fig. 1E). These results indicate a crucial role for both ANP32A&B in viral RNA replication.

**FIG 1.**
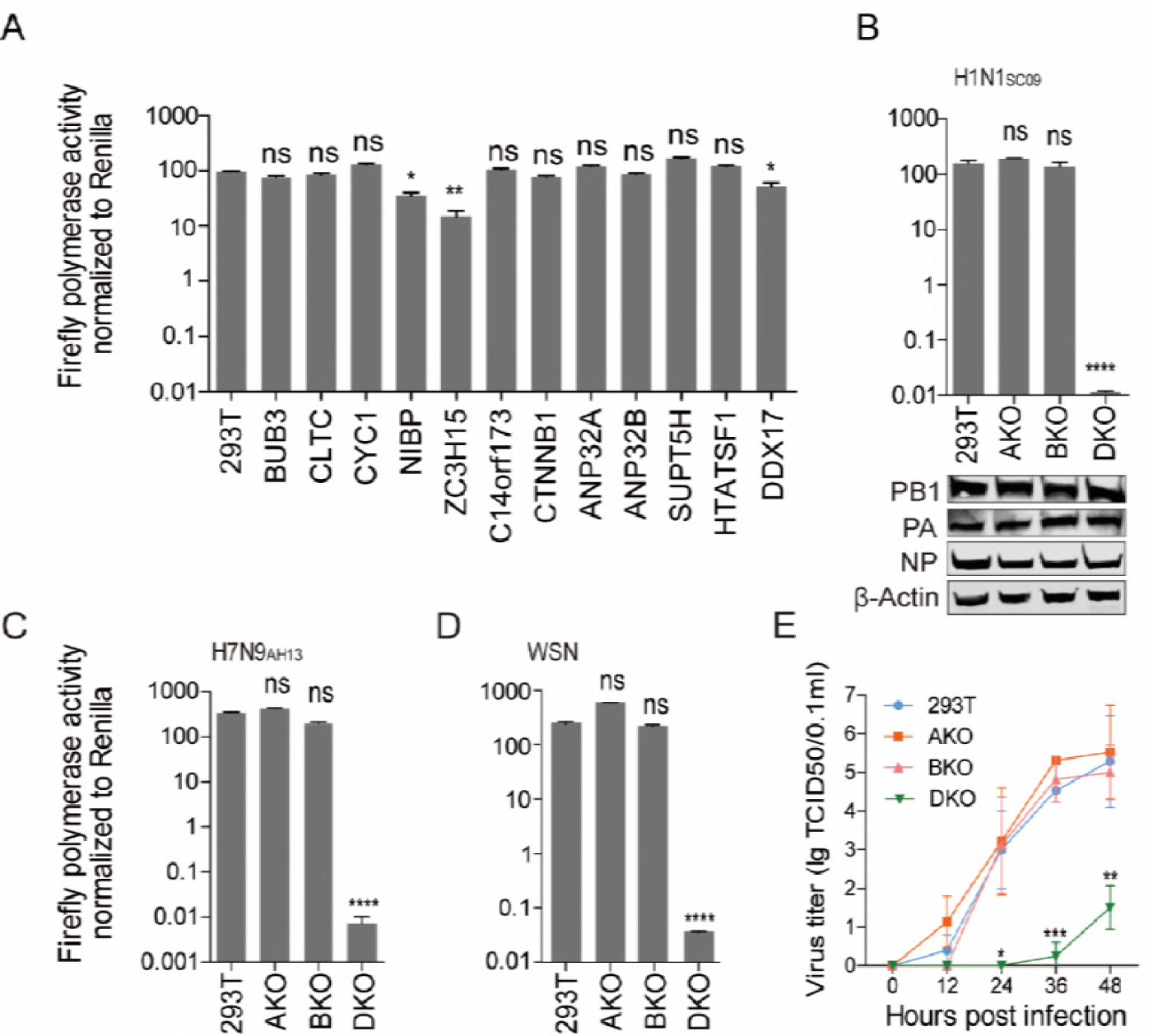
HuANP32A&B are indispensable for influenza A virus polymerase activity and viral replication. **(A)** Wild type 293T cells and single gene knockout 293T cell lines, including BUB3, CLTC, CYC1, NIBP, ZC3H15, C14orf173, CTNNB1, ANP32A, ANP32B, SUPT5H, HTATSF1, and DDX17, were transfected with firefly minigenome reporter, *Renilla* expression control, and the polymerase of H1N1_SC09_. Luciferase activity was measured 24 h after transfection and data are firefly Luciferase gene activity normalized to *Renilla* Luciferase gene activity. (Statistical differences between samples are indicated, according to a one-way ANOVA followed by a Dunnett’s test; NS= not significant, *P < 0.05, **P < 0.01, ****P < 0.0001. Error bars represent the SEM within one representative experiment.). (**B to D)** 293T, huANP32A knockout cells (AKO), huANP32B knockout cells (BKO), and huANP32A&B double knockout cells (DKO), were transfected with firefly minigenome reporter, *Renilla* expression control, and either H1N1_SC09_ polymerase (**B**), H7N9_AH13_ polymerase (**C**), or WSN polymerase (**D**). Luciferase activity was measured at 24 h post transfection. (**B** to **D,** data are firefly Luciferase gene activity normalized to that of *Renilla*, Statistical differences between cells are indicated, following a one-way ANOVA and subsequent Dunnett’s test; NS = not significant, **P < 0.01, ***P < 0.001, ****P < 0.0001. Error bars represent the SEM of the replicates within one representative experiment.). The expression levels of H1N1_SC09_ polymerase proteins were assessed by western blotting **(B)**. (**E)** 293T, AKO, BKO and DKO cells were infected with WSN virus at an MOI of 0.01. The supernatants were sampled at 0, 12, 24, 36, and 48 h post infection and the virus titers were determined by means of endpoint titration in MDCK cells. Error bars indicate SD from three independent experiments. *P < 0.05, **P < 0.01, ***P < 0.001.

The knockout of both ANP32A and ANP32B led to dramatic loss of viral polymerase activity (about 10000 fold), which is distinct from the previous report, which used a gene knockdown method and resulted in about 3-5 fold reduction of viral polymerase activity (8), indicating that ANP32A and ANP32B are of fundamental importance in viral replication. We found that reconstitution of either ANP32A or ANP32B, or both of them, in the DKO cells restored the viral polymerase activities of H1N1_SC09_, H7N9_AH13_, and WSN viruses (Fig. 2A to 2C). Expression of huANP32A or huANP32B in DKO cells supported the H1N1_SC09_ viral polymerase activity in a dose-dependent manner (Fig. 2D). Reconstitution of huANP32A or huANP32B or both of them in DKO cells restored full viral infectivity (Fig. 2E). We confirmed that the required level of expression of huANP32A or B is very low, and overdose expression has a negative effect (Fig. S2), suggesting an explanation of the confusing phenotype previously observed, that in normal 293T cells overexpression of ANP32 protein decreased viral polymerase activity (8). These data indicate that ANP32A and ANP32B are key factors required for polymerase activity, that they have similar functions in the support of viral replication, and that they can function independently and contribute equally to influenza virus polymerase activity.

**FIG 2.**
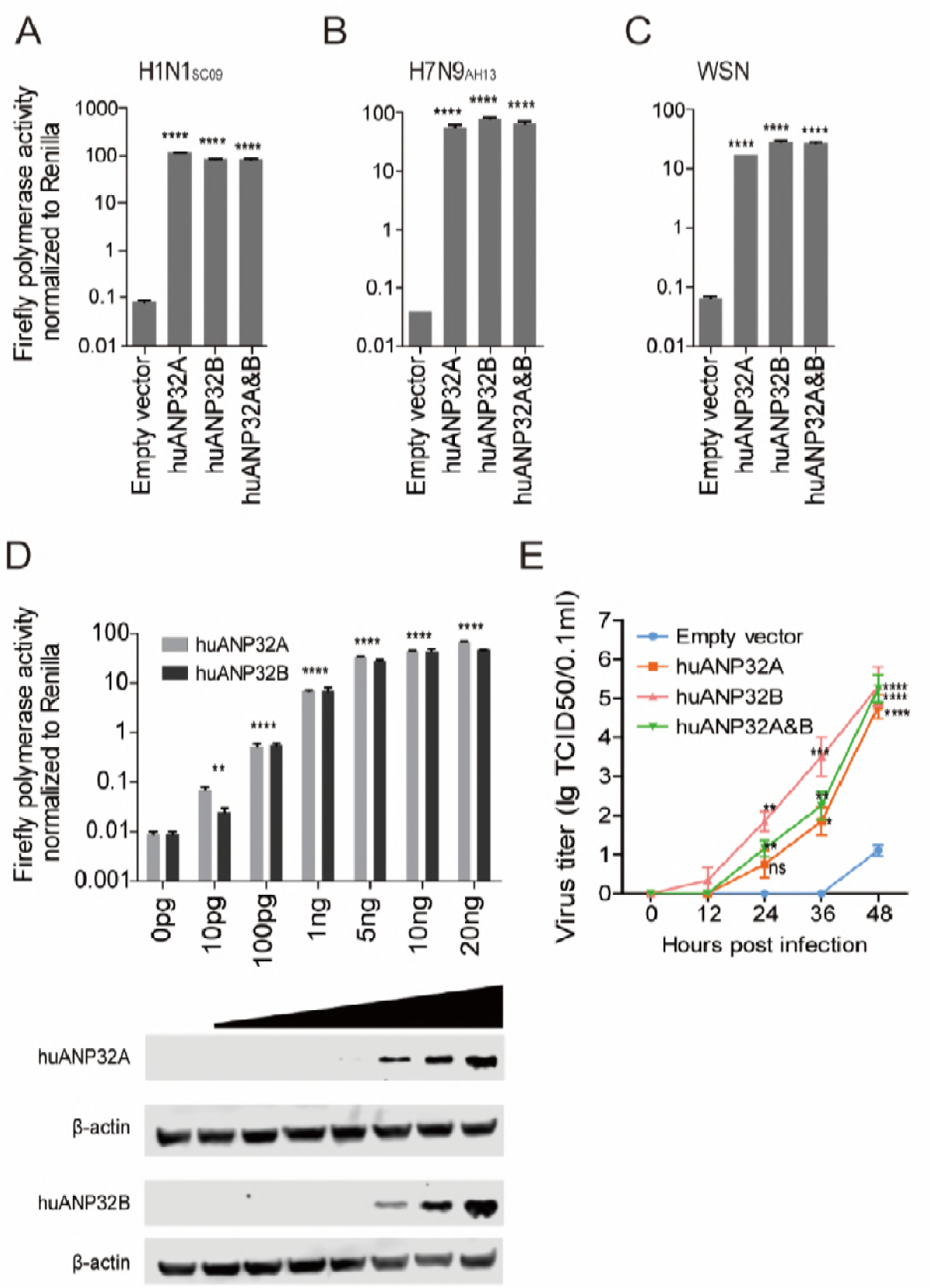
Reconstitutions of huANP32A&B rescue polymerase activity and replication of influenza A virus. 20 ng huANP32A, 20 ng huANP32B,10 ng huANP32A and 10 ng huANP32B, or 20 ng empty vector were co-transfected with either H1N1_SC09_ polymerase (**A**), H7N9_AH13_ polymerase (**B**) or WSN polymerase (**C**) into DKO cells, luciferase activity was assayed at 24 h after transfection. (**D)** Increasing doses of huANP32A or huANP32B were co-transfected with H1N1_SC09_ polymerase into DKO cells. Luciferase activity was measured at 24 h post transfection. The expression of ANP32 proteins was assessed by western blotting. (**A** to **D,** data are firefly Luciferase gene activity normalized to that of *Renilla*, Statistical differences between cells are indicated, following a one-way ANOVA and subsequent Dunnett’s test; NS = not significant, **P < 0.01, ***P < 0.001, ****P < 0.0001. Error bars represent the SEM of the replicates within one representative experiment). (**E**) DKO cells were transfected with 1ug huANP32A or/and B, or empty vector. After 24 h, the cells were infected with WSN virus at an MOI of 0.01. The supernatants were sampled at 0, 12, 24, 36, and 48 h post infection and the virus titers in these supernatants were determined as above. Error bars indicate SD from three independent experiments. NS = not significant, *P < 0.05, **P < 0.01, ***P < 0.001, ****P < 0.0001.

Influenza A virus replication starts from the transcription and replication of the negative single-stranded viral RNA (vRNA) by the vRNP complex. vRNA is transcribed into positive complementary RNA (cRNA) and mRNA, then the cRNA is used as a template to amplify into new vRNA (40, 41). We found that vRNA, cRNA, and mRNA synthesis were dramatically reduced in the DKO cells. However, when huANP32A was reconstituted in the DKO cells, vRNA, cRNA, and mRNA synthesis fully recovered (Fig. 3A to 3C). We observed similar results from reconstitution of ANP32B or both (Fig. 3A to 3C), indicating that huANP32A&B are key factors in triggering the replication of the viral genome.

**FIG 3.**
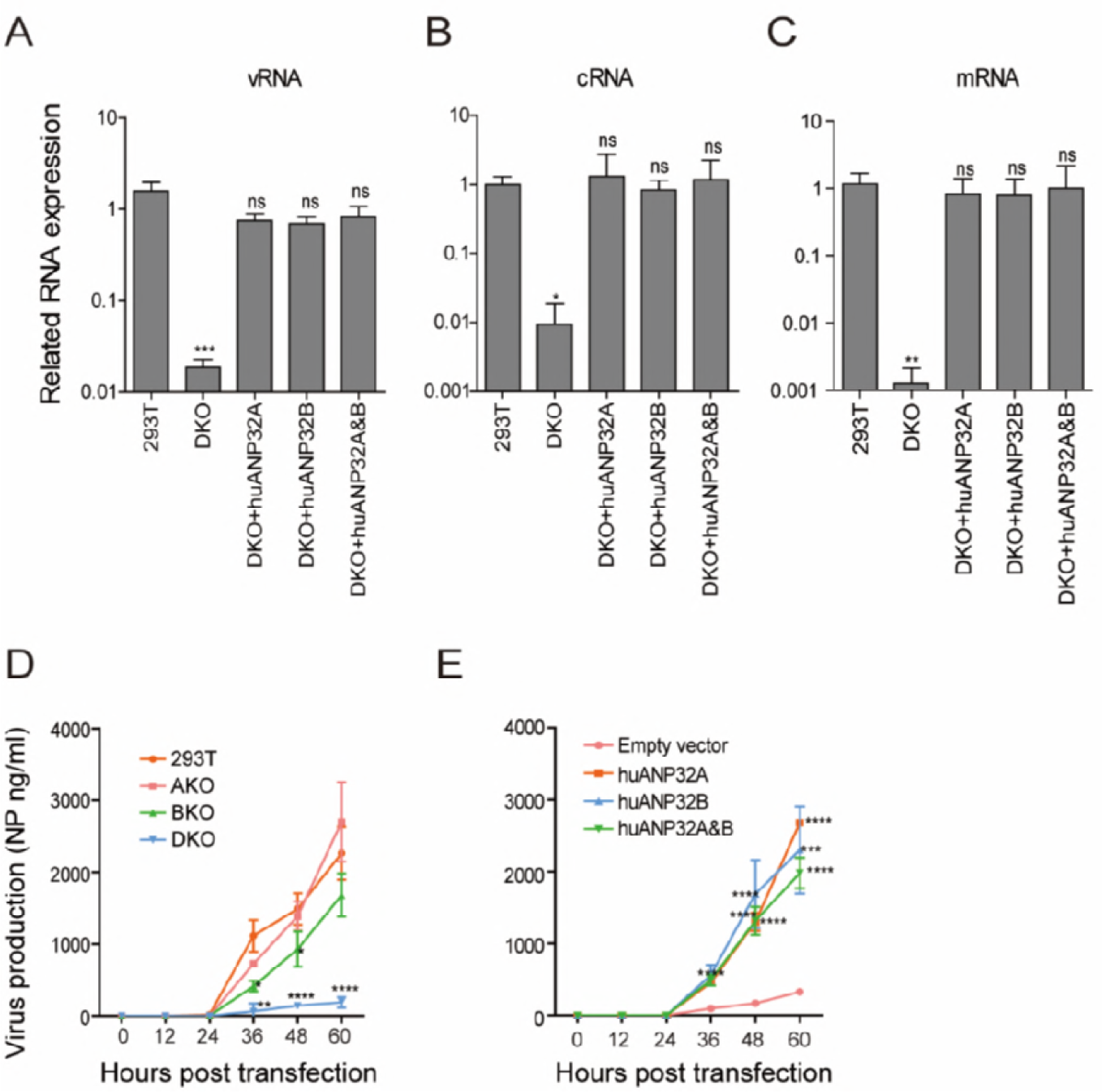
HuANP32A&B determine viral RNA replication efficiency and viral production. **(A-C)** 293T, or DKO cells, were transfected with the H1N1_SC09_ minigenome reporting system together with either 20 ng empty vector or huANP32A. The cells were incubated at 37 °C for 24 h before reverse transcription followed by quantitative PCR (qRT–PCR) for vRNA, mRNA and cRNA of the Luciferase gene. Error bars represent the SD from three independent experiments. NS = not significant, ****P < 0.0001. **(D** and **E)** Replication kinetics of H1N1_SC09_ in ANP32 knockout cells. 293T, AKO, BKO, and DKO cells in 6-well plates were transfected with 0.5 μg each of the eight plasmids of H1N1_SC09_ (**D**), or, in DKO cells, 40 ng huANP32A, 40ng huANP32B, 20 ng each of huANP32A and huANP32B, or 40 ng empty vector were co-transfected with the eight-plasmids of H1N1_SC09_ (**E**). The cells were cultured at 37 °C and supernatants collected at 0, 12, 24, 48 and 60 h post-transfection and subjected to virus production assay by ELISA, as described in ‘Methods’. Error bars represent the SD from three independent experiments. Error bars indicate SD from three independent experiments. *P < 0.05, **P < 0.01, ***P < 0.001, ****P < 0.0001.

We next used an eight-plasmid reverse genetics system in H1N1_SC09_ and tested the virus production in the supernatant of the transfected cells using an antigen capture ELISA (42). The result showed that the DKO cells had very low NP production in the cell supernatant, compared with the 293T, AKO, or BKO cells (Fig. 3D). The NP production was recovered when the huANP32A and/or huANP32B was expressed in the DKO cells (Fig. 3E). Taken together, these results suggest that huANP32A and huANP32B determine viral RNA replication efficiency and subsequent viral production in 293T cells.

### Support of Influenza A Viral Replication by ANP32A or B from Different Species

ANP32A&B are members of the evolutionarily conserved ANP32 family, which has various functions in the regulation of gene expression, intracellular transportat, and cell death (36). We tried to knock out ANP32A&B in both chicken DF1 cells and human A549 cells, but the knockout caused cell death, indicating both ANP32A&B have crucial roles in cell proliferation in these cells. ANP32A&B exist in almost all eukaryotic cells, with the exception of early eukaryotic life (yeast and other fungi) (43). Molecular phylogenetic analysis of ANP32 proteins revealed an evolutionary relationship among different species (Fig. S3). We then investigated the support of ANP32A or ANP32B from different species for viral polymerase activity in DKO cells. ANP32A from human, chicken, duck, turkey, zebra finch, mouse, pig, and horse, and ANP32B from human and chicken, were expressed individually with minigenomes of either H1N1_SC09_, human isolate H7N9_AH13_, H3N2 Canine influenza virus A/canine/Guangdong/1/2011(H3N2_GD11_), H3N8 equine influenza virus A/equine/Xinjiang/1/2007(H3N8_XJ07_), A/equine/Jilin/1/1989(H3N8_JL89_), or H9N2 chicken virus A/chicken/Zhejiang/B2013/2012(H9N2_ZJ12_), in DKO cells (Fig 4 and Fig. S4A). We found interestingly that chicken and other avian ANP32A proteins, which contain an additional 33 amino acids compared to the ANP32 proteins of mammals (8), supported viral polymerase activities in all cases, whereas the ANP32A proteins from human, pig, and horse, as well as human ANP32B, supported mammalian influenza virus polymerase activities, but not that of chicken H9N2_ZJ12_, or the H3N8_JL89_ (an avian-like virus). This result is consistent with previous reports that avian ANP32A can promote avian viral polymerase activity in human cells (8, 35). PB2 is the most important polymerase subunit, and affects host range (25, 44). Almost all the avian viruses had a glutamic acid (E) at PB2 residue 627, while it could be rapidly selected as a lysine (K) when the virus adapted to mammals, accompanied with increased pathogenicity and transmission abilities. The PB2 E627K mutation has long been regarded as a key signature of the avian influenza virus in overcoming the block to replicate in mammalian cells (9, 20, 21, 45-49). We observed that the K627E mutation in H7N9_AH13_ lost support from mammalian ANP32 proteins (Fig. S4A and S4B), while E627K mutations in isolates of avian origin (horse H3N8_JL89_, or chicken H9N2_ZJ12_) dramatically changed viral fitness to mammalian ANP32 proteins (Fig. S4B and S4C). These results revealed that ANP32A&B play important roles in influenza A viral replication across different species, and that mammalian ANP32 proteins provide poor support for avian influenza viral RNA replication.

**FIG 4.**
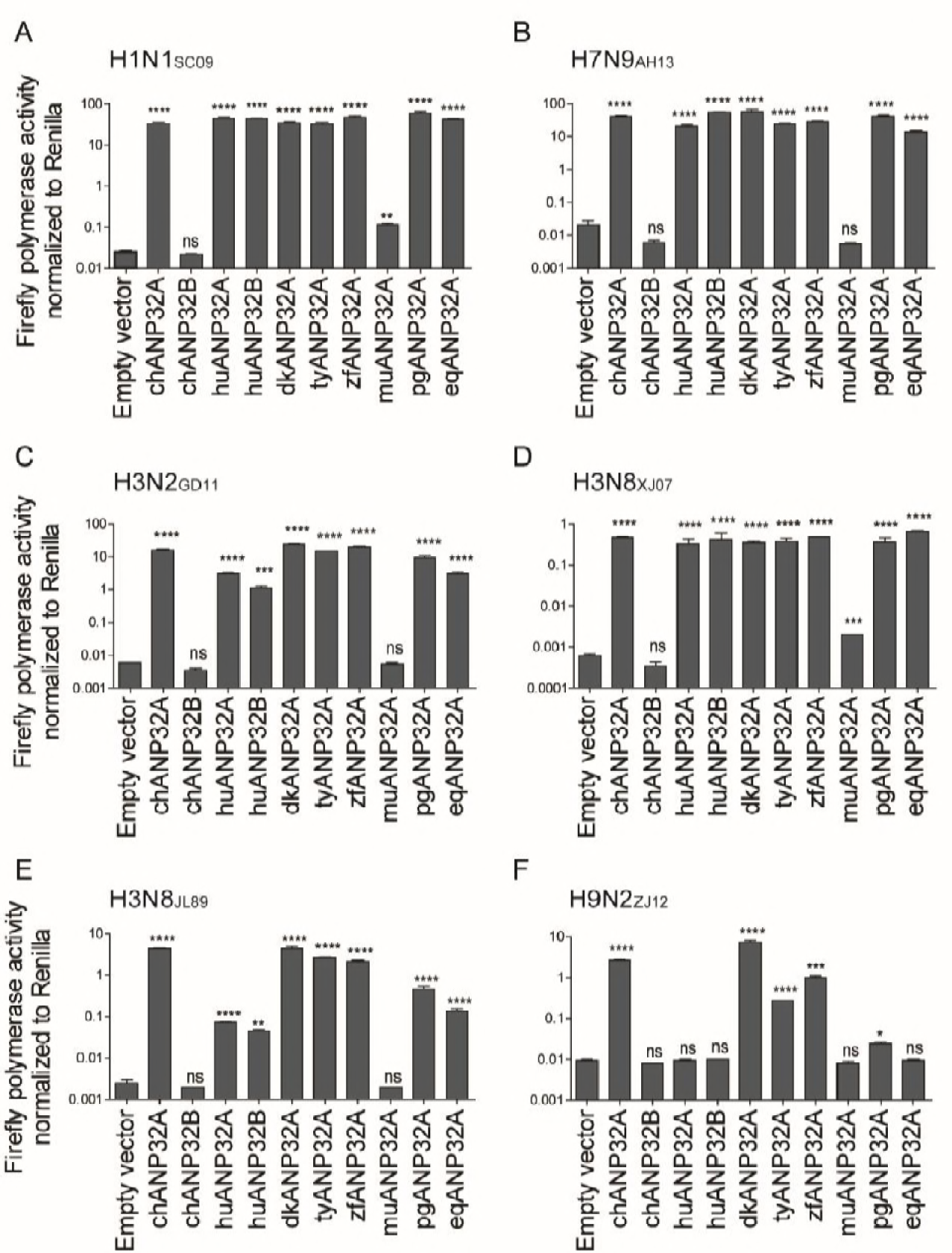
Support of influenza A viral replication by ANP32A or B from different species. Twenty nanograms of either empty vector or ANP32A or B from one of several different species was co-transfected with a minigenome reporter, *Renilla* expression control, and human influenza virus polymerase from H1N1_SC09_ (**A**), H7N9_AH13_ (**B**), canine influenza virus H3N2_GD11_ (**C**), equine influenza virus H3N8_XJ07_ (**D**) and H3N8_JL89_ (**E**), or avian influenza virus H9N2_ZJ12_ (**F**) into DKO cells. Luciferase activity was measured 24 h later. (Data are firefly Luciferase gene activity normalized to that of *Renilla*, Statistical differences between cells are indicated, following a one-way ANOVA and subsequent Dunnett’s test; NS = not significant, *P < 0.05, **P < 0.01, ***P < 0.001, ****P < 0.0001. Error bars represent the SEM of the replicates within one representative experiment.) dk, duck; ty, turkey; zf, zebra finch; mu, mouse; pg, pig; eq, equine.

### A Novel 129-130 Site Vital for Influenza Polymerase Activity in ANP32A&B

Surprisingly, chicken ANP32B did not support any viral RNA replication, and mouse ANP32A showed limited support to H1N1_SC09_ and H3N8_XJ07_, indicating a functional loss in these molecules (Fig. 4). The ANP32 protein family has a conserved structure containing an N-terminal leucine-rich repeat (LRR) and a C-terminal low-complexity acidic region (LCAR) domain (36). ANP32A&B have been reported to interact directly with the polymerase complex (7, 35). The key functional domains of ANP32 proteins remain largely unknown. We noticed that the chicken ANP32B (chANP32B) was functionally inactive and hypothesized that certain amino acids may be responsible for this phenotype. Alignment of huANP32B, pgANP32B, and chANP32B revealed scattered substitutions on the proteins (Fig. S5A). Chimeric clones between chicken and human ANP32B were constructed and evaluated (Fig. S5B). We found that the replacement of amino acids 111-160 of huANP32B with those of chANP32B aborted the activity of this protein, while chANP32B with human fragment 111-161 gained the ability to boost viral polymerase activity (Fig. S5C and S5D). Further comparing of ANP32B fragments 111-161 of chicken, human, and pig showed eight amino acid substitutions (Fig. 5A). Of these, a combined mutation N129I/D130N, but not others, on huANP32B impaired H1N1 polymerase activity (Fig. 5B), and the N129I showed stronger impairment than did the D130N mutation (Fig. 5C). We confirmed this phenotype on H7N9 polymerase activity (Fig. 5D). The 129-130 mutations also aborted the support of huANP32A of H1N1 and H7N9 polymerases (Fig. 5E and 5F). Furthermore, substitutions at 129-130 in chANP32A and chANP32B reversed the support of these proteins of the H7N9 virus human isolate (Fig. 5G). Interestingly, in the content of a chicken H7N9 viral minigenome, although mutation of 129-130 of chANP32A impaired the polymerase activity, the chANP32B reverse mutation did not restore its support to chicken-like H7N9 viral RNA replication (Fig. 5H), indicating that chicken virus replication may require both the 129-130 site and an extra 33-aminal acid peptide as in chANP32A. The expression levels of mutant ANP32 proteins were assessed using western blotting (Fig. S6). Most terrestrial mammalian ANP32A&B proteins have a 129-ND-130 signature, but most of the avian ANP32Bs have 129-IN-130 residues (Table S1). Murine ANP32A (muANP32A) has 129-NA-130 and this may explain the impaired support of H1N1 polymerase by muANP32A (Fig. 4 and Fig. S7). To further confirm the function of the 129-130 site, we made inactivating mutations of huANP32A and huANP32B at the sites 129-130 (N129I, D130N) on the 293T genome using CRISPR-Cas9 mediated recombination (Fig. 6A and Fig. S8); the double mutated proteins expressed well and fully impaired the replication of both H1N1_SC09_ and H7N9_AH13_ (Fig. 6B and 6C). Interestingly, the single ANP32A or ANP32B mutations lead to slightly decreased polymerase activities compared with wildtype 293T cells, suggesting competition between the mutated proteins and the native proteins for viral polymerase binding.

**FIG 5.**
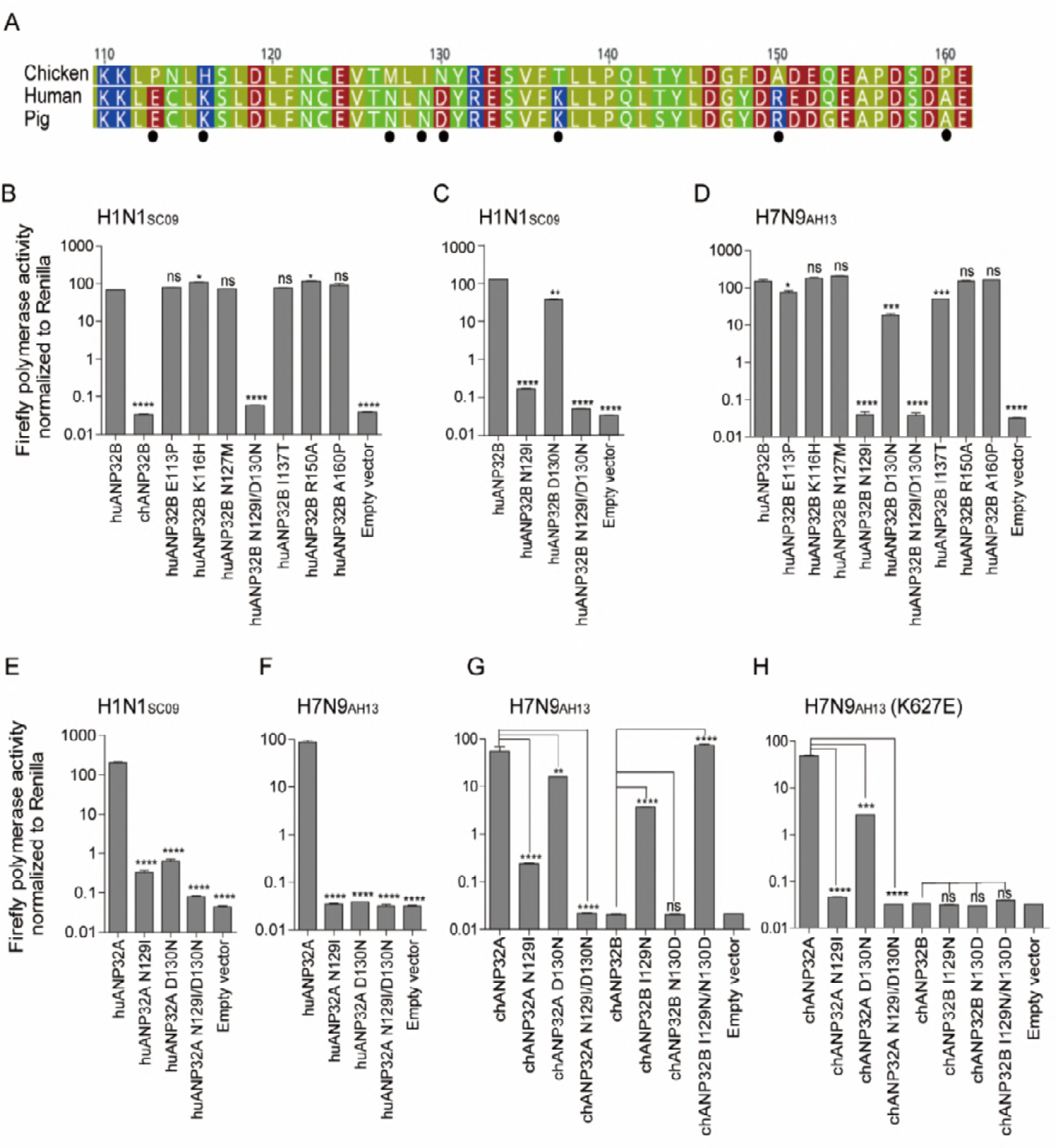
Key amino acids in ANP32A&B determine the activity of influenza viral polymerases. **(A)** Amino acid sequence comparison for chicken, human and pig ANP32B proteins (amino acids 110–161). Amino acid sequences are available in GenBank. The black dots indicate the different amino acid within these proteins. **(B** to **D)** huANP32B mutants were co-transfected with polymerase plasmids from H1N1_SC09_ (**B** and **C**), or H7N9_AH13_ (**D**), plus minigenome reporter and *Renilla* luciferase control into DKO cells. Luciferase activity was then assayed 24 h after transfection. **(E** and **F)** huANP32A mutants were co-transfected with polymerase plasmids from H1N1_SC09_ (**E**) or H7N9_AH13_ (**F**), plus a minigenome reporter and *Renilla* luciferase control, into DKO cells. **(G** and **H)** chANP32A or B mutants were co-transfected with polymerase plasmids from H7N9_AH13_ with either PB2 627K (**G**), or 627E (**H**) respectively. Luciferase activity was analyzed as mentioned above. (Data are firefly Luciferase gene activity normalized to that of *Renilla*, Statistical differences between cells are given, following one-way ANOVA and subsequent Dunnett’s test; NS = not significant, *P < 0.05, **P < 0.01, ***P < 0.001, ****P < 0.0001. Error bars represent the SEM of the replicates within one representative experiment.)

**FIG 6.**
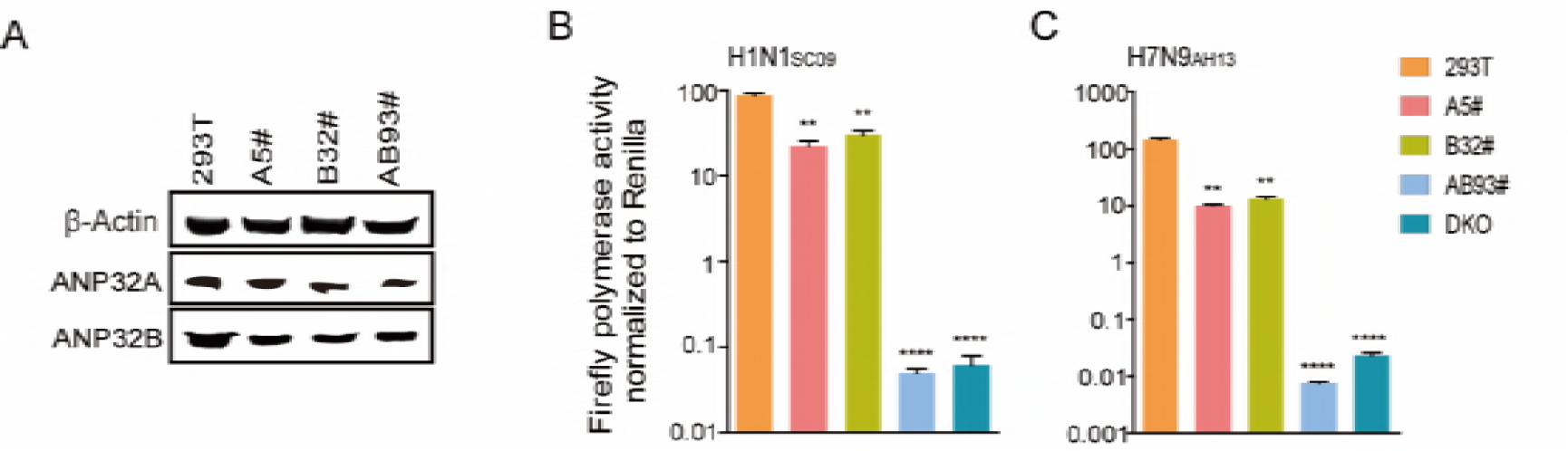
Amino-acids N129I/D130N substitute into ANP32A&B impaired influenza polymerase activity in 293T cells. (A) The endogenous protein expressions of different cell lines were identified by Western blotting using antibodies against β-actin, huANP32A and huANP32B. (B and C) Selected 293T cell lines were transfected with firefly minigenome reporter, *Renilla* expression control, and polymerase from H1N1_SC09_ (B) or H7N9_AH13_ (C). Cells were assayed for Luciferase activity. WT, wild-type 293T cells; A5#, ANP32A mutant cells; B32#, ANP32B mutant cells; AB93#, ANP32A/ANP32B double mutant cells. (Data are firefly Luciferase gene activity normalized to that of *Renilla*, Statistical differences between cells are given, following a one-way ANOVA and subsequent Dunnett’s test; NS = not significant, *P < 0.05, **P < 0.01, ***P < 0.001, ****P < 0.0001. Error bars represent the SEM of the replicates within one representative experiment.)

### The 129-130 Site in ANP32B Affects Its Interaction with Viral Polymerase

It has been reported that ANP32A variants interact with the polymerase subunit PB2 and that the avian ANP32A can enhance avian viral RNP assembly in human cells (35). ANP32A and ANP32B can interact with polymerase only when 3 subunits of the polymerase are present (7). ANP32B proteins from different species have a conserved structure that comprises a N-terminal leucine-rich repeat (LRR) region and a C terminal low complexity acidic region (LCAR) (Fig. 7A). To investigate the function of the 129-130 site in ANP32B in the interaction of this protein with viral polymerase, we used variants of huANP32B, chANP32B to co-transfect with WSN minireplicon into DKO cells. Co-immunoprecipitations detected strong interactions between huANP32A and the polymerase subunits PB1 and PA (Fig. 7B, Lane 1), but weak interactions when chANP32B was present (Fig. 7B, Lane 2). No interactions were detected when PB1 was absent (Fig. 7B, Lane 3), indicating this interaction occurred between ANP32B and the viral trimeric polymerase complex, which is consistent with previous findings (7, 34). As controls, we also observed that the LCAR truncations of huANP32B were unable to interact with the viral polymerase (Fig. 7B, Lane 4-6), in agreement with previous observations (34). When we mutated the functional human ANP32B at the site 129-130 to the chicken ANP32B signature, the huANP32B lost the ability to interact with viral polymerase. Conversely, chANP32B gained the ability to co-immumoprecipitate with viral polymerase when its 129-130 sites were mutated to the human ANP32B signature (Fig. 7C). Together these results revealed a fundamental function of ANP32A and ANP32B in supporting influenza A viral polymerase activity and a novel 129-130 site of ANP32A&B in different species that may influence influenza virus replication.

**FIG 7.**
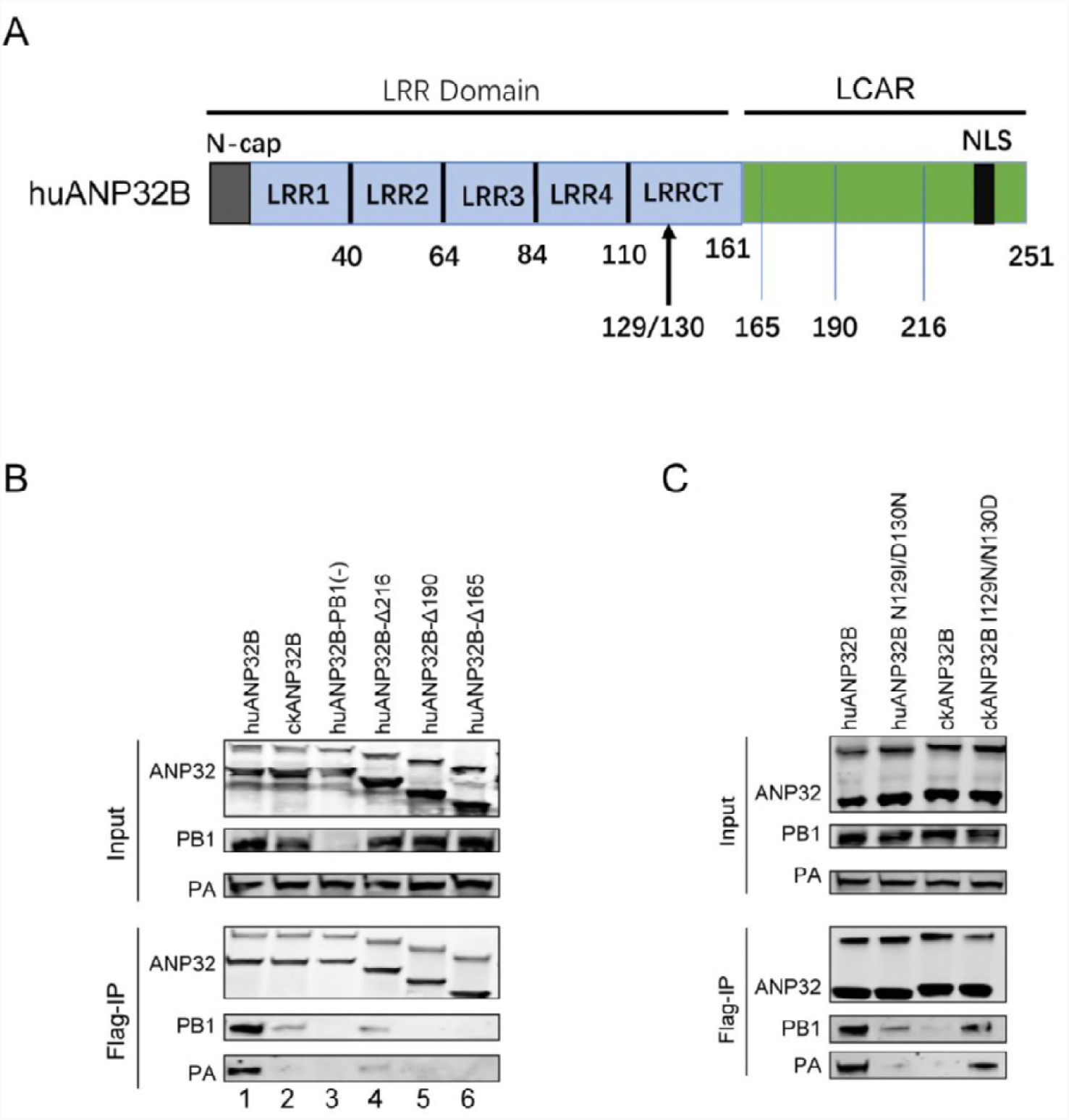
The 129-130 site in ANP32B contributes to the interaction with influenza polymerase. (A) A schematic describing the structure of ANP32B. LRR, leucine-rich repeat, including 4 LRR repeats and one C-terminal LRR domain (LRRCT); LCAR, low-complexity acidic region; NLS, nuclear localization signal. (B) and (C) 293T cells were transfected with different ANP32-Flag constructs, together with viral polymerase subunits PA, PB1, and PB2. Coimmunoprecipitation of the anti-Flag antibodies and the proteins was assessed using western blotting.

## DISCUSSION

Although the host factors that are involved in influenza A virus replication have been long investigated, and genome-wide screening has shown that many host proteins interact with viral polymerase, the key mechanisms that determine viral polymerase activity in different cells remain largely unclear. ANP32A and ANP32B have been previously identified binding with viral polymerase and promoting human influenza viral vRNA synthesis (7). Furthermore, a recent study showed for the first time that chicken ANP32A can rescue avian polymerase activity in human cells because the chicken ANP32A harbors an extra stretch of 33 amino acids which is absent in mammalian ANP32A (8). However, the potencies of ANP32A and ANP32B (as well as other host factors) in viral replication are not well investigated in different hosts. In our study, we used CRISPR/Cas9 knockout cells to screen the candidate host proteins involved in viral RNA replication. We identified that ANP32A and ANP32B are key host co-factors that determine influenza A virus polymerase activity. Without ANP32A&B, the viral polymerase activity decreased by 10000 fold and the viral infectivity decreased by more than 100000 fold. In contrast, the DDX17 knockout cells showed about 3-fold decreased viral polymerase activity (Fig. 1A), which was in agreement with prior reports (23). We found that ANP32A or ANP32B can independently restore viral polymerase activity, indicating that both have a similar function in viral replication (7).

Influenza viruses and their hosts have a long co-evolution history. ANP32 family members are expressed in animal and plant cells, with both conserved LRR and LCAR domains (43). Three conserved ANP32 members (ANP32A, ANP32B, and ANP32E) in vertebrates were reported to have several functions in cells. We found that knockout of both ANP32A and ANP32B abolished the influenza viral replication, while the ANP32E may not be involved in viral replication. Interestingly, we found that the chicken ANP32A keeps the conserved function to support both avian and mammalian virus polymerase activity, while human ANP32A or B did not support avian viral replication. Thus, we demonstrate here that ANP32A and ANP32B are key host factors that play a fundamental role in influenza A viral RNA replication.

A recent study revealed that avian ANP32A has a hydrophobic SUMO interaction motif (SIM) in an extra 33 amino acid insert region which connects to host SUMOylation to specifically promote avian viral polymerase activity (34). However, this SIM-like domain is not present in all of the ANP32B proteins from different hosts, and neither is it found in mammalian ANP32A. We showed that most ANP32Bs from different species are functional as ANP32As in support of viral replication, but chANP32B is naturally unable to support polymerase activity (Fig. 4). We finally found that the amino acids 129I and 130N of chANP32B are responsible for the loss of this function. Sequence analysis and mutagenesis studies suggest that the 129-130 sites are important for maintaining the function of avian ANP32A and human ANP32A/B in viral replication (Fig. 5). The co-immunoprecipitation assay showed that mutations on these 129-130 sites changed the interaction efficiency between the ANP32 proteins and viral polymerase complex (Fig. 7). Mutating the 129-130 sites in chANP32B to the functional signature 129N and 130D enables chANP32B to support polymerase activity in human viruses, but not chicken viruses with PB2 627E, indicating that the ANP32B may undergo selection in two areas during co-evolution with the avian influenza virus: chANP32B does not harbor an extra insert as chANP32A and the 129-130 mutations. This result also suggests that avian ANP32A is the only protein from the avian ANP32 family to support avian viral replication.

Together, out data give new insights into the functions of ANP32A and ANP32B. The 129-130 substitution could be used as a novel target to modify the genome of animals to develop influenza-A-resistant transgenic chickens, or other animals, which will benefit the husbandry industry, as well as animal and human health. Further investigation into the molecular mechanisms that determine how ANP32 proteins work with the viral polymerase complex, for example, the structure of the ANP32 and vRNP complex, and the fitness of different virus subtypes would contribute to the understanding viral pathogenesis and host defense.

## MATERIALS AND METHODS

### Cells, Viruses, and Plasmids

Human embryonic kidney (293T, ATCC CRL-3216) and Madin-Darby canine kidney (MDCK, CCL-34™) cells were maintained in Dulbecco’s modified Eagle’s medium (DMEM, Hyclone) with 10% fetal bovine serum (FBS; Sigma), 1% penicillin and streptomycin (Gibco), and kept at 37°C with 5% CO_2_. Certain reagents were kindly provided by the following persons: polymerase plasmids of H1N1 human influenza A virus A/Sichuan/01/2009 (H1N1_SC09_), and H7N9 A/Anhui/01/2013 (H7N9_AH13_) by Dr. Hualan Chen; H1N1 human influenza virus A/WSN/1933 (WSN) by Dr Kawoaka; H3N2 canine influenza virus A/canine/Guangdong/1/2011 (H3N2_GD12_) by Dr Shoujun Li from China Southern Agriculture University; and H9N2 avian influenza virus A/chicken/Zhejiang/B2013/2012 (H9N2_ZJ12_) by Dr Zejun Li from Shanghai Veterinary Research Institute of CAAS. H3N8 equine influenza viruses A/equine/Jilin/1/1989 (H3N8_JL89_) and A/equine/Xinjiang/1/2007 (H3N8_XJ07_) were preserved in our lab. The reverse genetics system based on the pBD vector for the H1N1_SC09_ virus was established in Dr Hualan Chen’s lab. The pCAGGS plasmids of containing full length ANP32A isoforms of several species were generated by gene synthesis (Synbio technologies, China) according to the sequences deposited in GenBank, including chicken ANP32A (chANP32A, XM_413932.5, XP_413932.3), human ANP32A(huANP32A, NM_006305.3, NP_006296.1), zebra finch ANP32A (zfANP32A, XM_012568610.1, XP_012424064.1), duck ANP32A (dkANP32A, XM_005022967.1, XP_005023024.1), turkey ANP32A (tyANP32A, XM_010717616.1, XP_010715918.1), pig ANP32A (pgANP32A, XM_003121759.6, XP_003121807.3), mouse ANP32A (muANP32A, NM_009672.3, NP_033802.2), equine ANP32A (eqANP32A, XM_001495810.5, XP_001495860.2), chicken ANP32B (chANP32B, NM_001030934.1, NP_001026105.1) and human ANP32B (huANP32B, NM_006401.2,m NP_006392.1). Site-directed mutants of these sequences were generated using overlapping PCR and identified by DNA sequencing. Mutants of pcAGGS-huANP32B-Δ216/190/165 and pcDNA3.1-PA-V5 were constructed according to the online In-Fusion ®HD Cloning Kit User Manual (http://www.clontech.com/CN/Products/Cloning_and_Competent_Cells/Cloning_Kits/xxclt_searchResults.jsp). Briefly, the fragments of the pcAGGS/pcDNA3.1 vector and each target gene were amplified with a 15 bp homologous arm and were then fused using the In-Fusion HD Enzyme (Clontech, Felicia, CA, USA). To create the pcAGGS-huANP32B-Δ216/190/165 plasmids, pcAGGS-huANP32B was used as the template to amplify the pcAGGS vector. This sequence was then fused with different truncated huANP32B fragments (huANP32B-Δ216/190/165). To obtain pcDNA3.1-PA-V5 plasmid, pBD-H1N1_SC09-_PA was used as the template to amplify the PA-V5 sequence, and then fused with pcDNA3.1 vector.

### Knockout Cell Lines

To generate knockout cell lines for host proteins BUB3(AF081496.1), CLTC(NM_004859.3), CYC1 (CR541674.1), NIBP (BC006206.2), ZC3H15 (NM_018471.2), C14orf173 (DQ395340.1), CTNNB1 (NM_001904.3), ANP32A (NM_006305.3), ANP32B (NM_006401.2), SUPT5H (U56402.1), HTATSF1 (NM_014500), and DDX17 (NM_006386, NM_001098504) (4, 23, 37), the gRNAs design tool (http://crispr.mit.edu/) was used for gRNAs design and off-target prediction (50). DNA fragments that contained the U6 promoter, gRNAs specific for host factors, a guide RNA scaffold, and U6 termination signal sequence were synthesized and subcloned into the pMD18-T backbone vector. The Cas9-eGFP expression plasmid (pMJ920) was a gift from Jennifer Doudna (Addgene plasmid # 42234) (51). Briefly, 293T cells in 6-well plates were transfected with 1.0 μg pMJ920 plasmids and 1.0 μg gRNA expression plasmids by Lipofectamine® 2000 Transfection Reagent (Invitrogen, Cat.11668-027) using the recommended protocols. GFP-positive cells were sorting by fluorescence-activated cell sorting (FACS) at 48 h post-transfection, then monoclonal knockout cell lines were screened by western blotting and/or DNA sequencing.

### Generation of Site-directed Amino-acid-substituted 293T Cell Line

High efficiency guide sequences for ANP32A and ANP32B which bind upstream and downstream with close proximity to the target (129/130 ND) were chosen. The gRNA expression plasmids were constructed as described above. An 80nt oligo with the desired mutations at the target site was used as the donor DNA. 293T cells were transfected with 1 μg pMJ920 plasmids, 1 μg gRNA expression plasmids, and 50 pmol donor DNA. 48 h later, GFP positive cells were isolated by FACS, and site-directed mutagenesis clones were identified after screening by sequencing and western blotting using Anti-PHAP1 antibody (ab51013) and Anti-PHAPI2 / APRIL antibody EPR14588 (ab200836). Cell lines with double gene mutations were generated by second round transfection and selection.

### Polymerase Assay

A minigenome reporter, which contains the firefly luciferase gene flanked by the non-coding regions of the influenza HA gene segment with a human polI promoter and a mouse terminator sequence (52), was transfected with viral polymerase and NP expression plasmids to analyze the polymerase activity. Mutants of PB2 genes were generated using overlapping PCR and identified by DNA sequencing. To determine the effect of ANP32 proteins on viral polymerase activity, 293T or KO cells in 12-well plates were transfected with plasmids of the PB1 (80 ng), PB2 (80 ng), PA (40 ng) and NP (160 ng), together with 80 ng minigenome reporter and 10 ng *Renilla* luciferase expression plasmids (pRL-TK, kindly provided by Dr Luban), using Lipofectamine 2000 transfection reagent (Invitrogen) according to the manufacturers’ instructions. Cells were incubated at 37°C. Cells were lysed with 100ul of passive lysis buffer (Promega) at 24 h after transfection, and firefly and *Renilla* luciferase activities was measured using a Dual-luciferase kit (Promega) with Centro XS LB 960 luminometer (Berthold technologies). The function of ANP32 was examined using a polymerase assay by co-transfection of DKO cells with different ANP32 proteins for 24 h. All the experiments were performed independently at least three times. Results represent the mean ± SEM of the replicates within one representative experiment. The expression levels of polymerase proteins on different cell lines were detected by western blotting, using specific mouse monoclonal antibodies for NP and PB1 proteins, anti-V5 tag antibody (ab27671) for PA-V5 protein.

### RNA isolation, Reverse Transcription, and Quantification by RT-PCR

Total RNA from 293T cells was extracted using an RNeasy mini kit (Qiagen), following the manufacturer’s instructions. For the synthesis of first strand cDNA derived from firefly luciferase RNAs driven by influenza polymerase, equal concentrations of RNA were subject to cDNA synthesis using a Reverse Transcription Kit (PrimeScript™ RT reagent Kit with a gDNA Eraser (Perfect Real Time), Cat.RR047A). Primers used in the RT reaction were as follows: 5’-CATTTCGCAGCCTACCGTGGTGTT-3’ for the firefly luciferase vRNA; 5’-AGTAGAAACAAGGGTG-3’ for the firefly luciferase cRNA; oligo-dT20 for the firefly luciferase mRNA (53). The cDNA samples were subjected to quantification by real-time PCR with specific primers F (5’-GATTACCAGGGATTTCAGTCG-3’) and R (5’-GACACCTTTAGGCAGACCAG-3’) using SYBR ® Premix Ex TaqTM II (Tli RNaseH Plus) (TaKaRa Cat: RR820A). Fold change in RNA was calculated by double-standard curve methods and β-actin as an internal control.

### Quantitative ELISA for Determination of Virus Production

The ELISA assay has been previously described(42). Briefly, a 96-well microtiter plate (Costar, Bodenheim, Germany) was coated with 1μg/well of the mouse monoclonal anti-IAV NP protein antibody in phosphate-buffered saline (PBS), incubated overnight at 4 °C or 37 °C for 2 h. The plate was washed three times with washing buffer (PBS containing 0.1% Tween-20, PBST) and blocked with 200 μl 5% calf serum at 37 °C for 2 h. After washing three times with PBST buffer, 100 μl of virus/VLPS-containing samples were added in dilution buffer (PBS containing 10% calf serum and 0.1% Triton X-100) and incubated at 37 °C for 1 h. The plate was then washed, and 100 μl of a 1:2000 dilution of HRP-conjugated anti-IAV NP protein mAb was added. After incubation at 37 °C for 0.5 h, the plate was washed again and incubated with freshly prepared TMB peroxidase substrate (Galaxy Bio, Beijing, China) for 10 min at room temperature. The reaction was stopped by adding 2 M H_2_SO_4_, and the optical density at 450 nm wavelength (OD450) was measured using the VersaMax Microplate Reader (BioTek, Winooski, USA). Dilution buffer was taken as a blank control and the purified IAV-NP protein was double diluted eight times in dilution buffer to obtain the standard curve. The virus production was calculated using the standard curve.

### Influenza Virus Infection and Infectivity

Human influenza virus WSN was rescued from a 12-plasmid rescue system (54). Briefly, 293T culture in a 6-well plate was transfected with 0.1μgof each pPolI plasmids and 1ug each of pCAGGS-NP, pCAGGS-PA, pCAGGS-PB1, and pCAGGS-PB2 using Lipofectamine 2000 (Invitrogen) in Opti-MEM. Eight hours post-transfection, medium was changed to DMEM plus 10% FBS. The supernatant was harvested after 48 h and injected into 9-day-old specific pathogen-free embryonated eggs for virus propagation. Eggs were incubated at 35 °C for 72 h, the allantoic fluid from eggs was detected by haemagglutination assay and the titers of virus on 293T cells were determined using the method of Reed and Muench (55). Then 293T cells, huANP32A knockout 293T cells (AKO cells), huANP32B knockout 293T cells (BKO cells), and huANP32A &B double knocked out 293T cells (DKO) were infected at a multiplicity of infection (MOI) of 0.01 for 1 h, washed twice with PBS, and then cultured at 37°C in Opti-MEM containing 0.5% FBS and tosylsulfonyl phenylalanyl chloromethyl ketone (TPCK)-trypsin (Sigma) at 1mg/ml. At the indicated time points, the culture supernatant was harvested. Viral titers in MDCK cells were determined as described above.

### Immunoprecipitation and Western Blotting

For immunoprecipitation and western blotting, transfected cells were lysed using an ice-cold lysis buffer (50 mM Hepes-NaOH [pH 7.9], 100 mM NaCl, 50 mM KCl, 0.25% NP-40, and 1 mM DTT), and centrifuged at 13,000× g and 4 °C for 10 min. After centrifugation, the crude lysates were incubated with Anti-FLAG M2 Magnetic Beads (SIGMA-ALDRICH, M8823) or Anti-NP Magnetic Beads (MCE Protein A/G Magnetic Beads and NP antibody from our lab) at 4°C for 2 hr. After incubation, the resins were collected using a magnetic separator and washed three times with PBS. The resin-bound materials were eluted using a 3X Flag peptide or by boiling in the SDS-PAGE loading buffer, subjected to SDS-PAGE and then transferred onto nitrocellulose membranes. Membranes were blocked with 5% milk powder in Tris-buffered saline (TBS) for 2 h. Incubation with the first anti-mouse antibody (Anti-Flag antibody from SIGMA (F1804), Anti-V5 antibody from Abcam (ab27671), Anti-NP and PB1 antibody from our lab) was performed for 2 h at room temperature (RT), followed by washing three times with TBST. The secondary antibody (Sigma, 1:10,000) was then applied and samples were incubated at RT for 1 h. Subsequently, membranes were washed three times for 10 min with TBST. Signals were detected using a LI-COR Odyssey Imaging System (LI-COR, Lincoln, NE, USA).

### Statistics

Statistical analysis was performed in GraphPad Prism, version 5 (Graph Pad Software, USA). Statistical differences between groups were assessed by One-way ANOVA followed by a Dunnett’s post-test. All the experiments were performed independently at least three times. Error bars represent the standard deviation (SD) or the standard error of the mean (SEM) in each group, as indicated in figure legends. NS, not significant (p>0.05), *p<0.05, **p<0.01, ***p<0.001, ****p<0.0001.

## ACKNOWLEDGEMENTS

We thank W. Barclay, SJ. Li, ZJ. Li, J. Luban, and J. Doudna for providing the plasmids and advice. We thank CJ. Li and DM. Zhao for discussions.

## SUPPORTING INFORMATION

**FIG S1.**
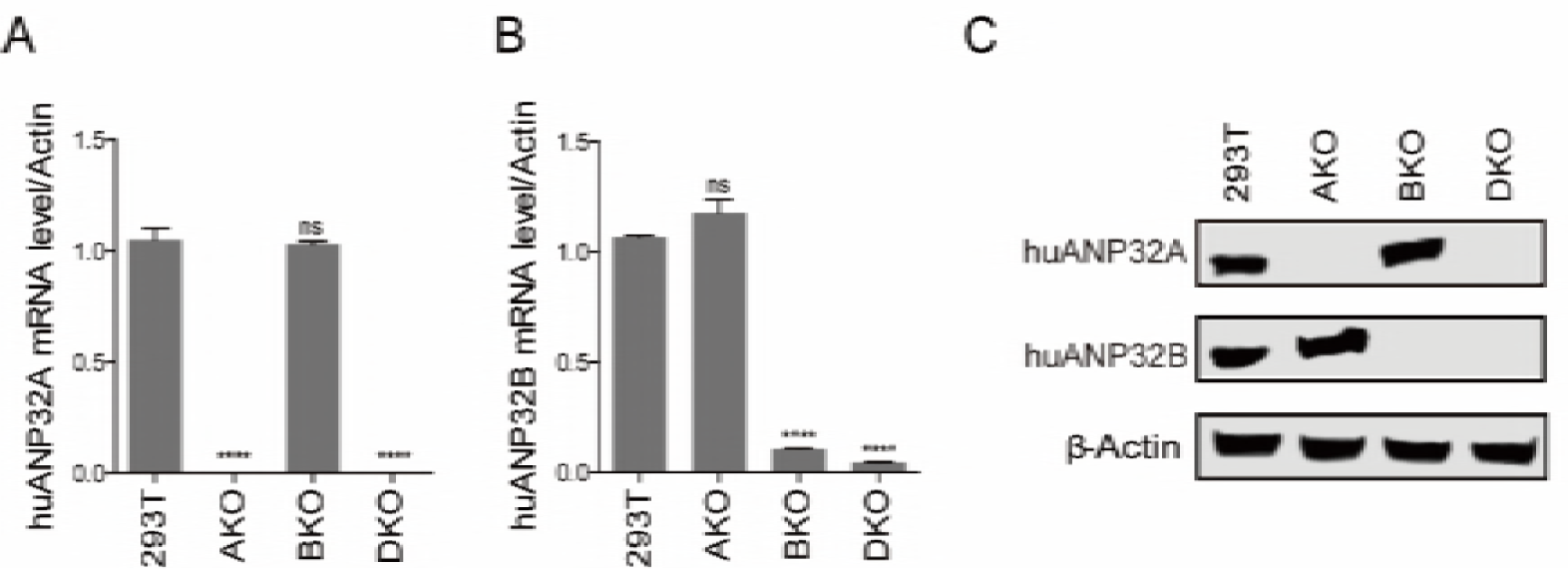
The identification of huANP32A and huANP32B knockout cells. 293T cells were transfected with pMJ920 vector (plasmid expressing eGFP and Cas9) and gRNAs targeting huANP32A or huANP32B to generate huANP32A knockout cells (AKO), huANP32B knockout cells (BKO), and huANP32A&B double knockout cells (DKO). The mRNA levels of huANP32A and B in different cell lines were determined using qRT-PCR using primers specific for huANP32A (**A**) and huANP32B (**B**). The primer sequences are given in ‘Materials and Methods’. Fold change in RNA was calculated by double-standard curve methods. β-actin was used as an internal control. Statistical differences between samples are given, following one-way ANOVA and subsequent Dunnett’s test; NS = not significant, ****P < 0.0001. Error bars indicate SD from three independent experiments. (**C**) The endogenous proteins were identified by western blotting using antibodies against β-actin, huANP32A and huANP32B, described as “Materials and Methods”.

**FIG S2.**
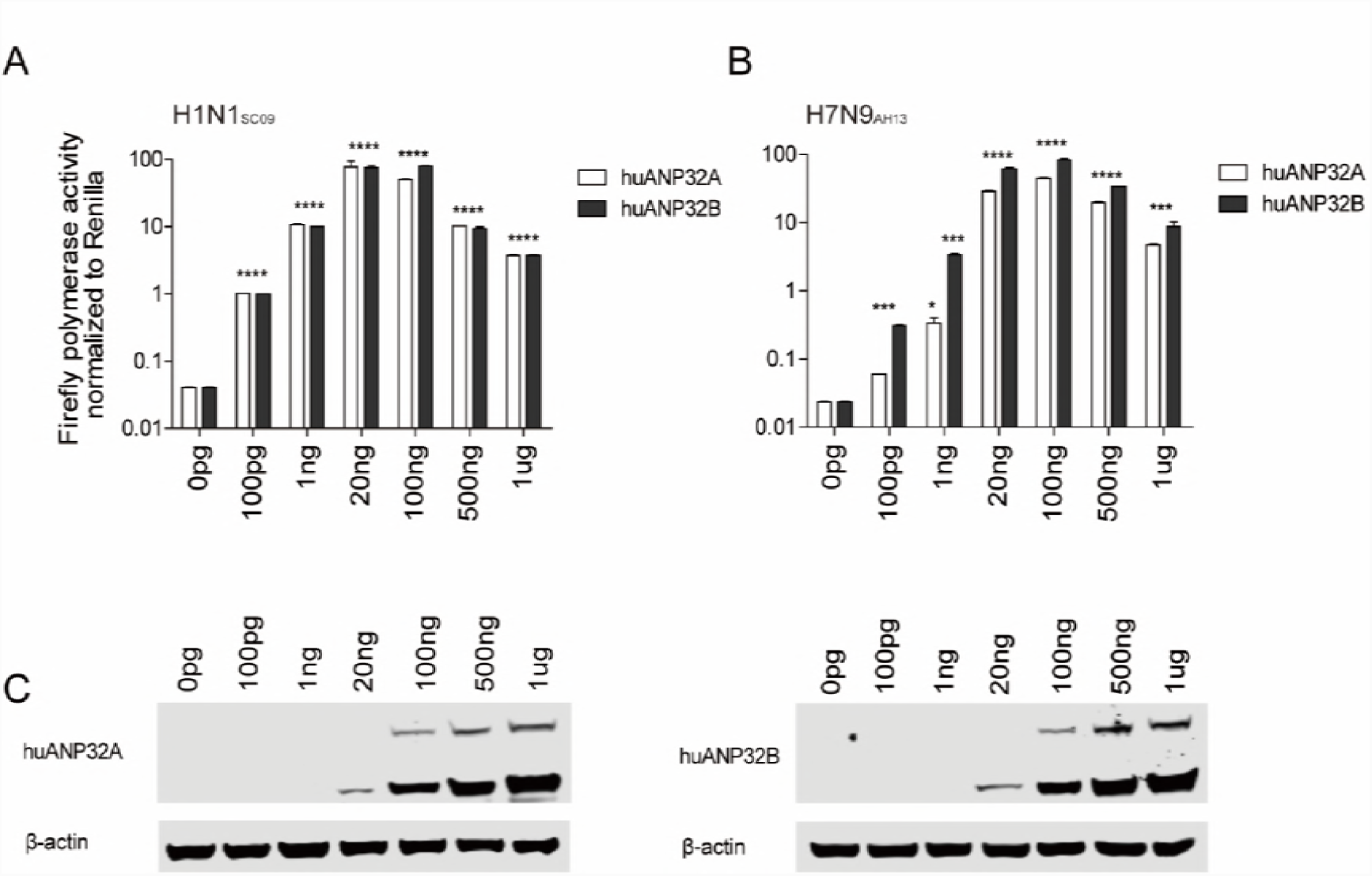
Overdose expression of huANP32A&B impaired viral polymerase activity. Different doses of huANP32A or huANP32B were co-transfected with H1N1_SC09_ polymerase (**A**) and H7N9_AH13_ polymerase (**B**) into DKO cells. Luciferase activity was measured 24 h after transfection and data are firefly Luciferase gene activity normalized to that of *Renilla*. (Statistical differences between samples are given, following one-way ANOVA and subsequent Dunnett’s test; NS = not significant, *P < 0.05, ***P < 0.001, ****P < 0.0001. Error bars represent the SEM of the replicates within one representative experiment.). The expression levels of the huANP32A and huANP32B proteins were confirmed by western blotting.

**FIG S3.**
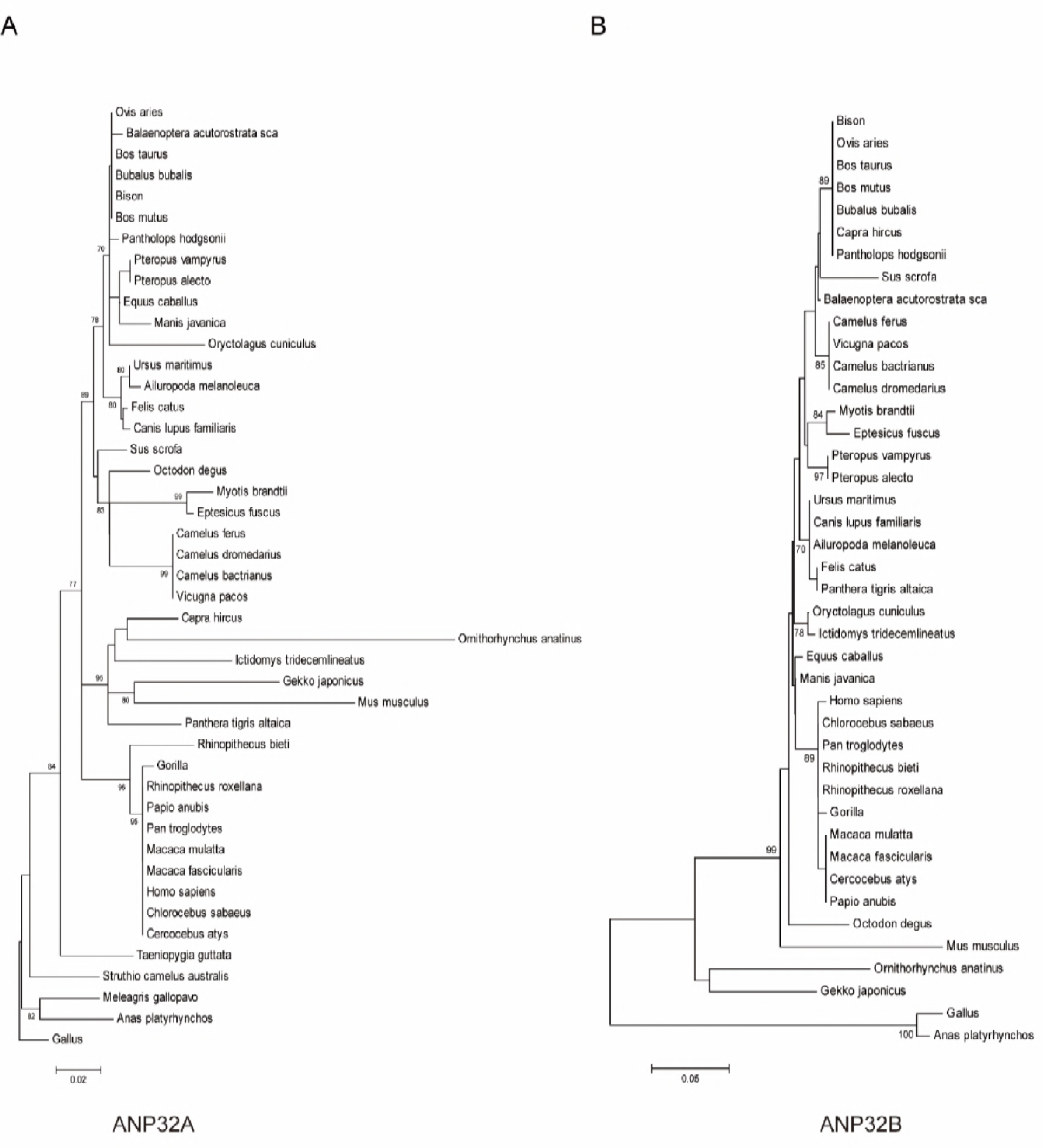
Phylogenetic tree of amino acid sequences of ANP32A (A) and ANP32B (B). Trees were generated by the neighbor-joining method and evaluated by 1000 bootstrap pseudo replicates. The bar scale represents the genetic distance. The credibility value for each node is shown.

**FIG S4.**
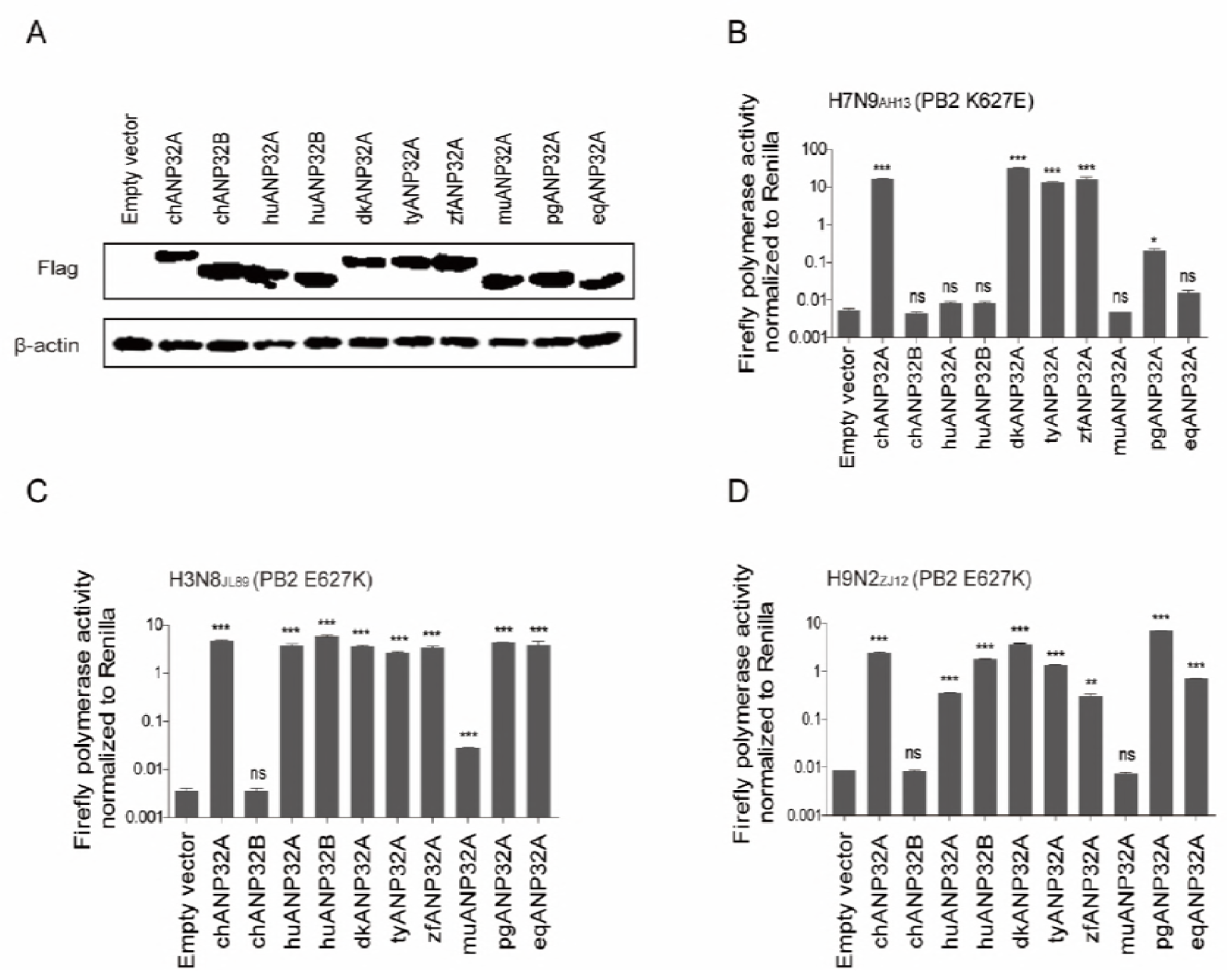
Expression of flag-tagged ANP32A or B proteins and PB2 E627K mutation overcomes human ANP32A&B restriction. (A) 1µg of each ANP32 plasmid was transfected into 293T cells using Lipofectamine® 2000. 48 h post transfection, the cell lysates were analyzed using SDS-PAGE and Western Blotting using antibodies against Flag peptide and β-actin. (B-D) DKO cells were transfected with 20ng either empty vector or ANP32A or B from one of several different species, together with minigenome reporter, *Renilla* expression control, and different viral polymerase including: human H7N9_AH13_ with PB2 K627E (**B**), an avian origin equine influenza virus H3N8_XJ07_ with PB2 E627K (**C**), and avian influenza virus H9N2_ZJ12_ with PB2 E627K (**D**). Luciferase activity was measured 24 h post transfection. (Data are polymerase activity normalized to *Renilla*; Statistical differences between cells are given, following one-way ANOVA and subsequent Dunnett’s test; NS = not significant, *P < 0.05, **P < 0.01, ***P < 0.001, ****P < 0.0001. Error bars represent the SEM of the replicates within one representative experiment.) dk, duck; ty, turkey; zf, zebra finch; mu, mouse; pg, pig; eq, equine.

**FIG S5.**
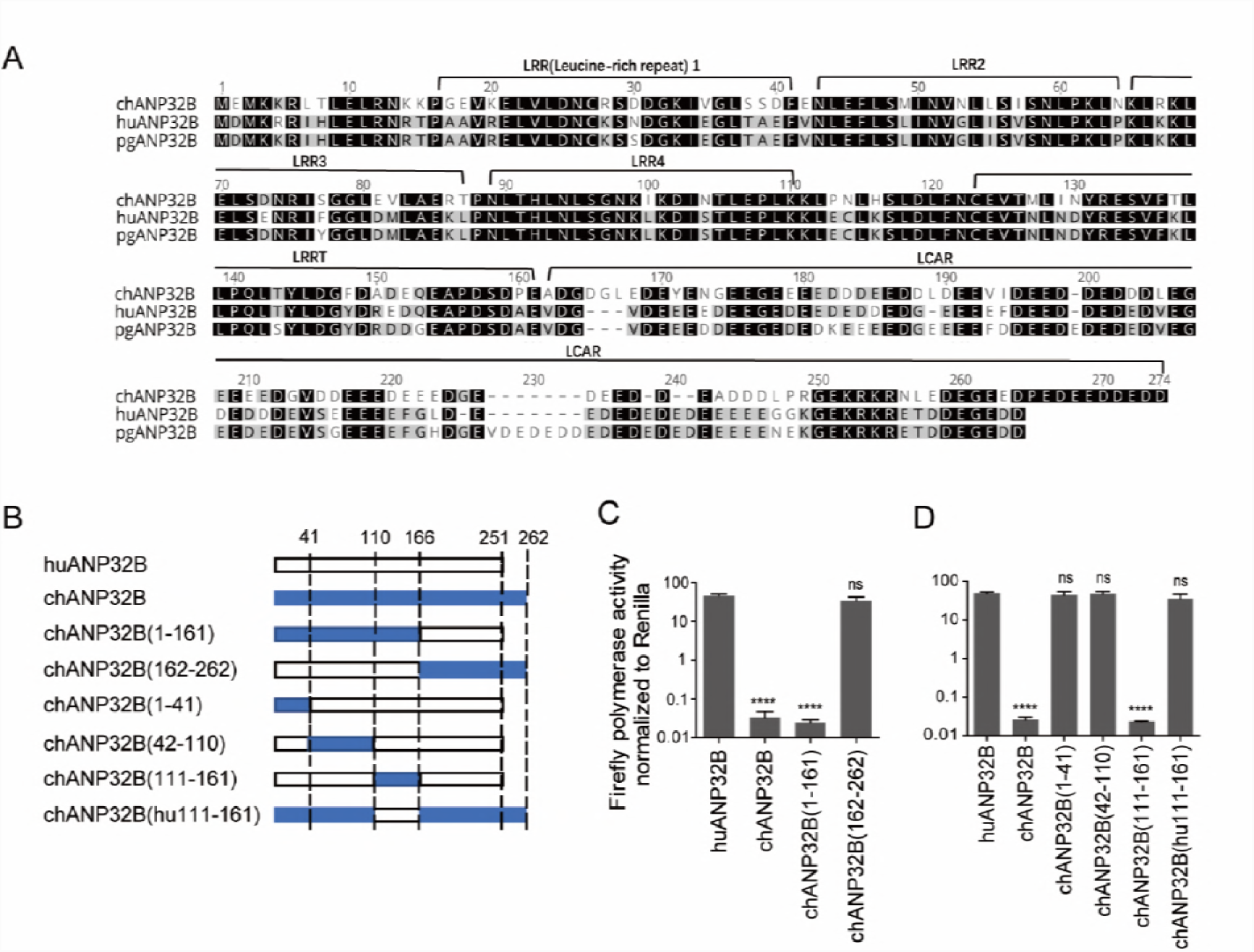
Mapping of crucial sites in ANP32 proteins that influence viral polymerase activity. (A) ANP32B sequences from chicken, human, and pig were aligned using the Geneious R6 software. The notches are marked with dashes. The similarity of amino acid identity is highlighted in different colors. (B) Schematic diagram of chimeric clones between chicken and human ANP32B, constructed according to the known domains (Separated by dotted lines.1-41aa, LRR1 region; 42-110aa, LRR2,3 &4 region; 111-161aa, LRRCT region; 162-251aa, LCAR region, as showed in a). The bars show the origin of the genes as follows: white, huANP32B; blue, chANP32B. (56) Chimeric clones were co-transfected with minigenome reporter, *Renilla* expression control, and H1N1_SC09_ polymerase into DKO cells. Luciferase activity measured 24 h later. (Data are polymerase activity normalized to *Renilla*; Statistical difference between samples are given labeled, following one-way ANOVA and subsequent Dunnett’s test; NS = not significant, ****P < 0.0001. Error bars represent the SEM of the replicates within one representative experiment.)

**FIG S6.**
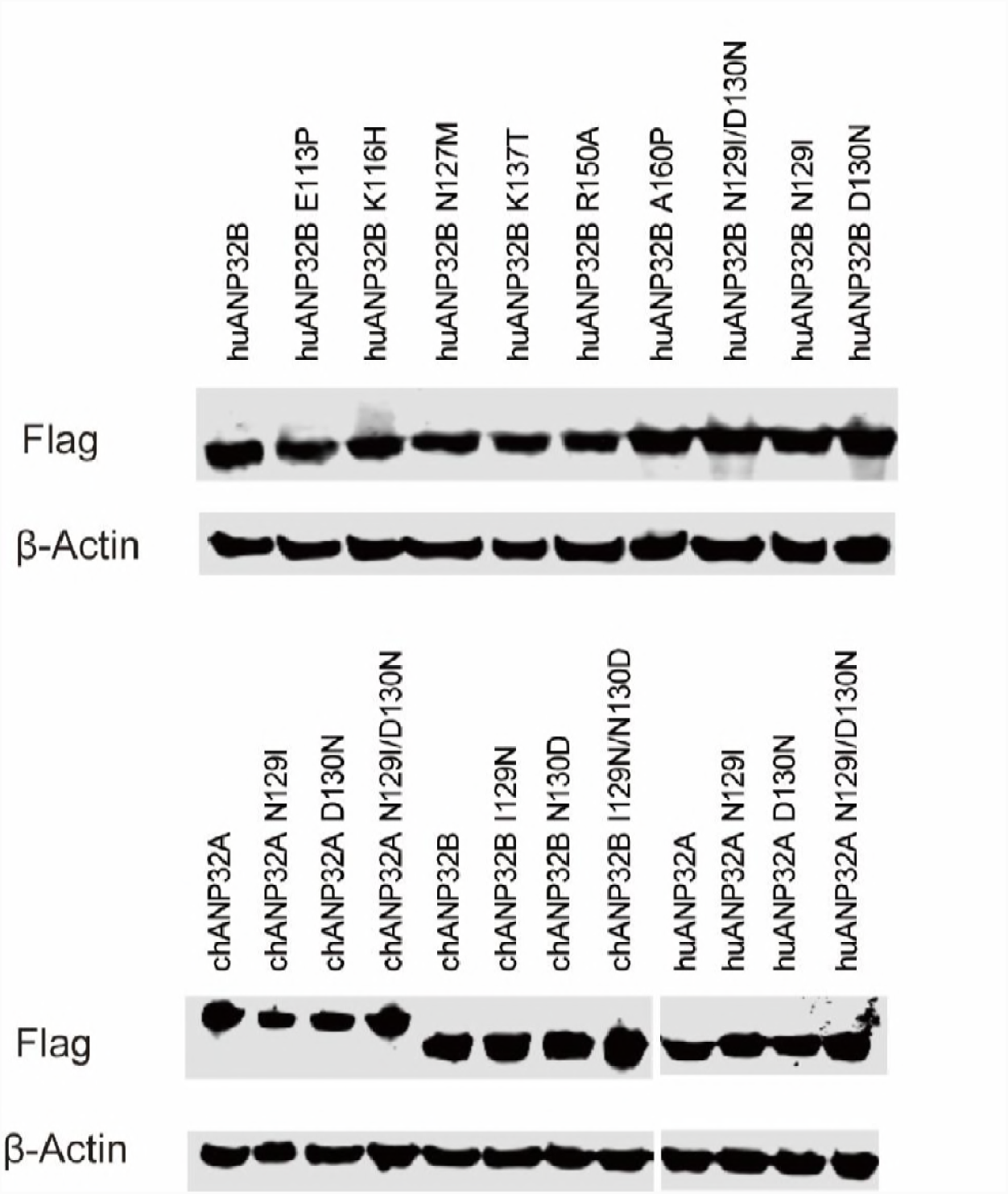
Expression of flag-tagged ANP32A or B mutants. 1µg of each mutant ANP32 and wildtype ANP32 plasmids were transfected in 293T cells by Lipofectamine® 2000. 48 h post transfection, the cells lysates were analyzed by SDS-PAGE and Western Blotting using antibodies against Flag peptide and β-actin.

**FIG S7.**
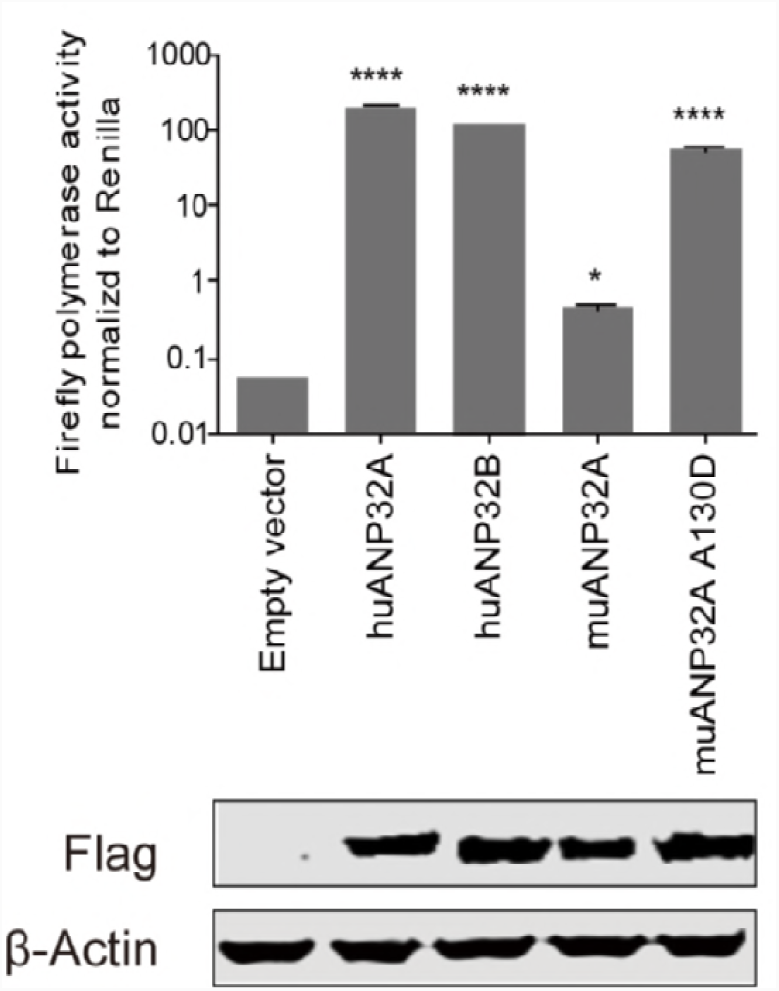
Mouse ANP32A 130A impaired its support of viral replication. Mutated muANP32A or human ANP32 proteins were co-transfected with minigenome reporter, *Renilla* expression control, and H1N1_SC09_ polymerase into DKO cells. Luciferase activity was assayed. (Data are polymerase activity normalized to *Renilla*; Statistical differences between cells are given, following one-way ANOVA and subsequent Dunnett’s test; NS = not significant, *P < 0.05, ****P < 0.0001. Error bars represent the SEM of the replicates within one representative experiment.)

**FIG S8.**
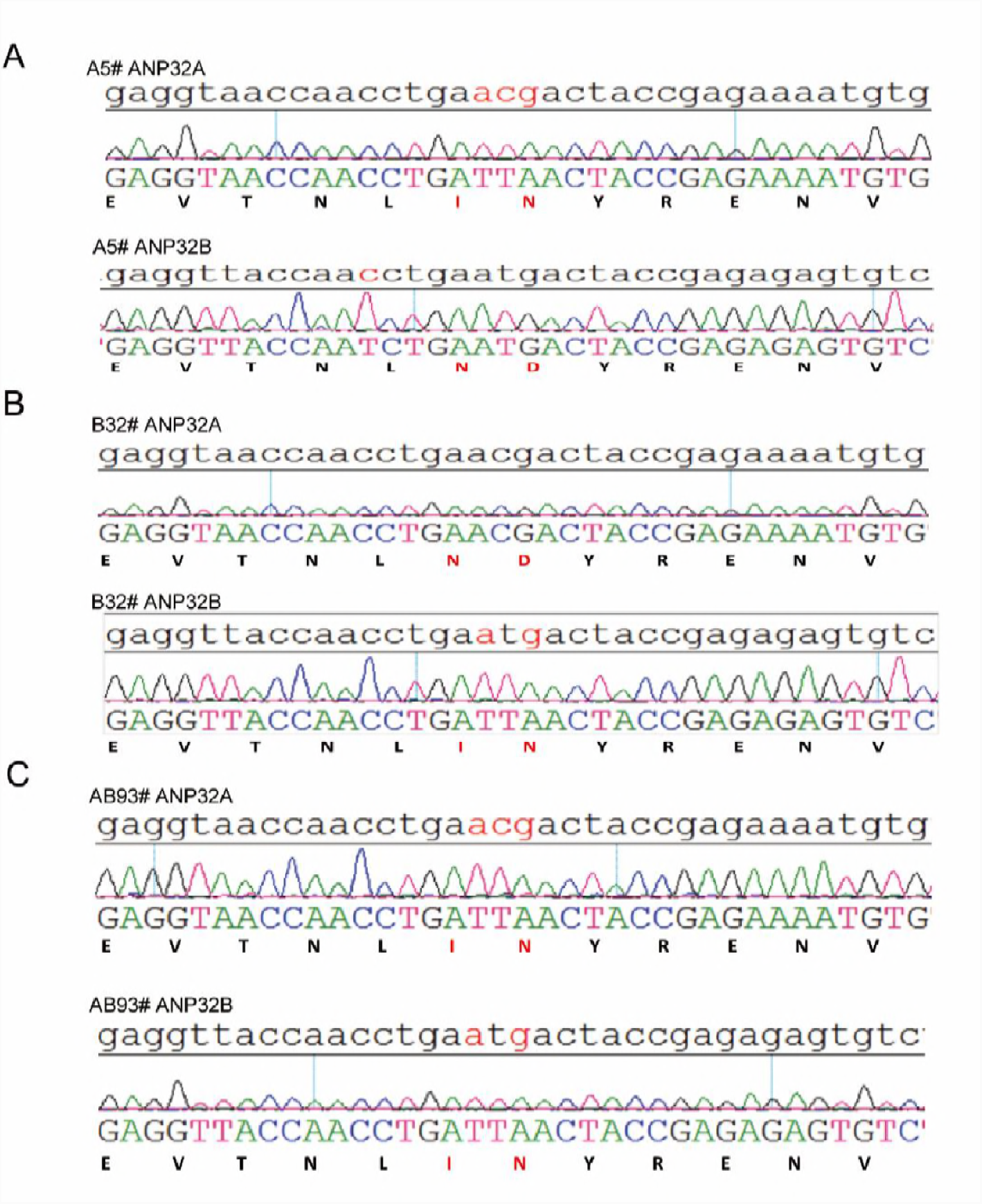
Identification of ANP32A&B site-mutation 293T cell lines. N129I/D130N substitutions in huANP32A and ANP32B on the 293T chromosome were generated using a CRISPR/Cas9 system. Positive 293T colonies of N129I/D130N substitutions on huANP32A and ANP32B were identified by sequencing. (A) Cloned cell A5# was identified carrying the N129I/D130N on ANP32A but not ANP32B. (B) Cell B32# was identified harboring N129I/D130N on ANP32B but not ANP32A. (C) Cell AB93# has N129I/D130N on both ANP32A&B.

**Table S1.** Summary of amino acids at 129-130 of ANP32 proteins

|  | Residue at amino acid position* |  |  |  |
| --- | --- | --- | --- | --- |
|  | ANP32A |  | ANP32B |  |
| Host | 129 | 130 | 129 | 130 |
| *Gallus* | N | D | I | N |
| *Anas platyrhynchos* | N | D | I | N |
| *Homo sapiens* | N | D | N | D |
| *Gorilla gorilla* | N | D | N | D |
| *Felis catus* | N | D | N | D |
| *Canis lupus familiaris* | N | D | N | D |
| *Bos taurus* | N | D | N | D |
| *Sus scrofa* | N | D | N | D |
| *Mus musculus* | N | A | S | D |
| *Equus caballus* | N | D | N | D |

931 \*The sequences of different species ANP32 proteins were retrieved from NCBI

